# No evidence that sex and transposable elements drive genome size variation in evening primroses

**DOI:** 10.1101/007161

**Authors:** J Arvid Ågren, Stephan Greiner, Marc TJ Johnson, Stephen I Wright

## Abstract

Genome size varies dramatically across species, but despite an abundance of attention there is little agreement on the relative contributions of selective and neutral processes in governing this variation. The rate of sexual reproduction can potentially play an important role in genome size evolution because of its effect on the efficacy of selection and transmission of transposable elements. Here, we used a phylogenetic comparative approach and whole genome sequencing to investigate the contribution of sex and transposable element content to genome size variation in the evening primrose (*Oenothera*) genus. We determined genome size using flow cytometry from 30 *Oenothera* species of varying reproductive system and find that variation in sexual/asexual reproduction cannot explain the almost two-fold variation in genome size. Moreover, using whole genome sequences of three species of varying genome sizes and reproductive system, we found that genome size was not associated with transposable element abundance; instead the larger genomes had a higher abundance of simple sequence repeats. Although it has long been clear that sexual reproduction may affect various aspects of genome evolution in general and transposable element evolution in particular, it does not appear to have played a major role in the evening primroses.

## Introduction

Variation in genome size is one of the most striking examples of biodiversity (Bennett and Leitch 2012; Gregory 2013). Genomes may be as small as 160 kb, as in the obligate endosymbiotic proteobacterium *Carsonella ruddii* (Nakabachi et al. 2006), and as large as 150 GB in the polyploid plant *Paris japonica* (Pellicer et al. 2010). Variation is not restricted to differences between species, as extensive genome size variation also exists within species (Biemont 2008; Diez et al. 2013; Long et al. 2013). Understanding the evolutionary processes underlying this variation has received much attention (reviewed in for example Petrov 2001; Gregory 2005; Lynch 2007; Gaut and Ross-Ibarra 2008; Ågren and Wright 2011), yet little consensus exists on the relative contributions of these processes.

Variation in genome size may be influenced by both neutral and selective evolutionary processes. Several studies have shown that genome size may evolve neutrally, with increases and decreases mainly due to biases in insertion and deletion rates and/or recombination rates (Petrov 2001; Oliver et al. 2007; Nam and Ellegren, 2012). It has also long been recognized that genome size correlates with various ecologically relevant traits; with examples ranging from flowering time in plants (Meagher and Vassiliadis, 2005) to powered flight in birds (Wright et al. 2014) and selection may thus act adaptively on genome size. Much recent debate has focused on the hypothesis that variation in the efficacy of selection, usually due to differences in effective population size (*N_e_*), governs most variation in genome size across distantly related taxa (see for example exchanges by Lynch and Conery 2003; Charlesworth and Barton 2004; Daubin and Moran 2004; Lynch and Conery 2004; Whitney et al. 2010, Whitney and Garland 2010; Lynch 2011; Whitney et al. 2011). Finally, several studies show that differential accumulation of transposable elements (TEs) can explain differences in genome size (reviewed in Ågren and Wright 2011). Indeed, there appears to be a correlation between relative TE abundance and genome size across angiosperms (Tenaillon et al. 2010). Furthermore, variation between closely related species, including species of rice (Piegu et al. 2006), cotton (Hawkins et al. 2006), and *Arabidopsis* (Hu et al. 2011), can be attributed to differences in TE abundance. Understanding what factors allow TEs to accumulate in some species, but not in others, may therefore be central to understanding variation in genome size in general, and among plants in particular.

A key factor affecting the efficacy of selection and TE transmission is the rate of sexual reproduction (Hickey 1982; Wright and Schoen 1999; Morgan 2001; Docking et al. 2006; Dolgin and Charlesworth 2006; Glemin and Galtier 2012; Ågren 2014). Sexual reproduction provides a way for TEs to spread to new lineages in a population, and despite the deleterious effects on host fitness, theory predicts that TEs in sexual populations should evolve maximum transposition rates (Charlesworth and Langley 1986). By contrast, within-lineage transmission of genes within asexual populations is expected to limit TE spread and in the absence of horizontal gene transfer reduction in transposition rate due to self-regulation is more likely to evolve in highly selfing and asexual species (Charlesworth and Langley 1986). Taken together, this leads to the prediction of higher TE abundances and larger genome sizes in sexually reproducing species compared to asexually reproducing relatives. On the other hand, the reduced effective population size in asexuals means that any actively transposing TEs they inherit from sexual progenitors may initially accumulate to high numbers by a Muller’s ratchet-like process (Muller 1964; Gabriel et al. 1993; Charlesworth 2012). Over time, asexual lineages with high TE abundance are expected to go extinct, whereas asexual lineages with low TE abundance are more likely to persist (Arkhipova and Meselson 2005; Dolgin and Charlesworth 2006). Whether sex will cause an increase or decrease in genome size remains unclear and represents an important problem for understanding the forces that affect genome evolution.

To date, most studies on the role of sexual reproduction on TE accumulation and genome expansion have focused on animal systems (e.g Zeyl et al. 1996; Arkhipova and Meselson 2000; Schaack et al. 2010) whereas work on plants has been lacking (but see Docking et al. 2006). Studies on plants have typically compared self-fertilizing species to outcrossing relatives (e.g Lockton and Gaut 2010; de la Chaux et al. 2012; Ågren et al. 2014). Several studies have shown an association between outcrossing rate and genome expansion (Albach and Greilhuber 2004; Trivers et al. 2004; Wright et al. 2008; Hu et al. 2011), but it is difficult to distinguish whether these differences are due to TE accumulation in the outcrossers or loss of TEs in the selfers (Wright and Ågren 2011). Furthermore, when correcting for phylogenetic non-independence, Whitney et al. (2010) failed to detect a significant effect of outcrossing rate on genome size in a broad scale comparison of 205 species. Similar large-scale studies of the effect of sexual reproduction on plant genome size evolution have so far been lacking.

The evening primrose plant family (Onagraceae) provides an ideal system to test the effect of sexual reproduction on genome size evolution. Functional asexuality (i.e. absence of recombination and segregation) in the Onagraceae has evolved more than 20 times independently due to a genetic system called Permanent Translocation Heterozygosity (PTH), which is characterized by suppression of meiotic recombination and segregation (Cleland 1972; Harte 1994; Rauwolf et al. 2008; Johnson et al. 2009). Functional asexuality due to PTH has been described in eight plant families (Cleland 1972; Holsinger and Ellstrand, 1984; Harte 1994), and differs from apomixis in that individuals go through all stages of meiosis, and successful zygote formation still requires fertilization (see Whitton et al. 2008 for a review of other forms of plant asexuality). Another useful contrast between PTH and many other asexual species is that PTH species tend to share the same ploidy level with their sexual relatives, allowing the effect of reproductive system to be decoupled from the effect of ploidy.

In this study we take a phylogenetic comparative approach to examine whether the repeated transitions from sex to PTH in *Oenothera* has been associated with a shift in genome size. We also use whole genome sequencing of three species of varying genome size and reproductive system to assess the contribution of transposable elements to genome size variation in the genus.

## Material and Methods

### Study system

In this study we focus on the *Oenothera* genus of the evening primrose family Onagraceae. The genus is monophyletic (Levin et al. 2004; Wagner et al. 2007) and includes the largest number of PTH species and their sexual relatives (Raven 1979; Holsinger and Ellstrand 1984; Johnson et al. 2011). PTH in *Oenothera* results from three mechanisms (Cleland 1972; Harte, 1994; Rauwolf et al. 2008). First, during metaphase I of meiosis, all 14 chromosomes form a ring (*x* = *n* = 7) instead of bivalents, which restricts synapsis and recombination to the highly homozygous telomeres, such that genetically detectable recombination is effectively 0. Second, in anaphase I, alternate disjunction results in one haploid set of chromosomes segregating as a unit, and the other haploid set as another unit. Unless one of each unit is present in the zygote, the zygote will not survive. This balanced mortality of gametes prevents segregation, which leads to permanent heterozygosity. Finally, > 99.5% of seeds are self-fertilized (R Godfrey and MTJ Johnson, unpublished results) because receptive stigmas accept pollen before flowers open. In short, the genetics of PTH reproduction in *Oenothera* can be likened with splitting the genome in half; only to later fuse the two halves back together, without recombination or segregation (Cleland 1972; Harte, 1994; Rauwolf et al. 2008). Recent evidence shows that recombination is also suppressed in sexual bivalent forming species, suggesting that recombination rates may not be dramatically different between sexual and PTH species. However, sexual species still undergo free segregation of homologous chromosomes, which should effectively eliminate genetic linkage disequilibria between chromosomes and allow the formation of homozygous loci from heterozygous parents as per Hardy-Weinberg expectations (Rauwolf et al. 2011; Golczyk et al. 2014). By contrast, these processes are completely lost in PTH species, leading to the perpetual propagation of single genotypes (Stebbins 1950; Cleland 1972).

### Genome size estimates

In this study we examined genome size in 30 *Oenothera* species, including 17 PTH and 13 sexual species (Figure 1). Sterilized seeds were stratified by sowing them on agar and kept in the dark at 4°C for three weeks. Seedlings were transferred to pots and grown in controlled glasshouse conditions between October 2013 and January 2014 at the University of Toronto. Leaf tissue, sampled at the same level of maturity (∼ 5 weeks), from each species was sent to Plant Cytometry Services (PO Box 299, 5480 AG Schijndel, The Netherlands), who determined DNA content in picograms (pg) using flow cytometry with Propidium Iodide fluorescent dye and *Pachysandra terminalis* as a standard (1C = 1.73; Zonneveld et al. 2005). This method has previously been shown to successfully detect small differences in genome size, including within species variation (Diez et al. 2013). Three replicates were performed per species.

**FIGURE 1.**
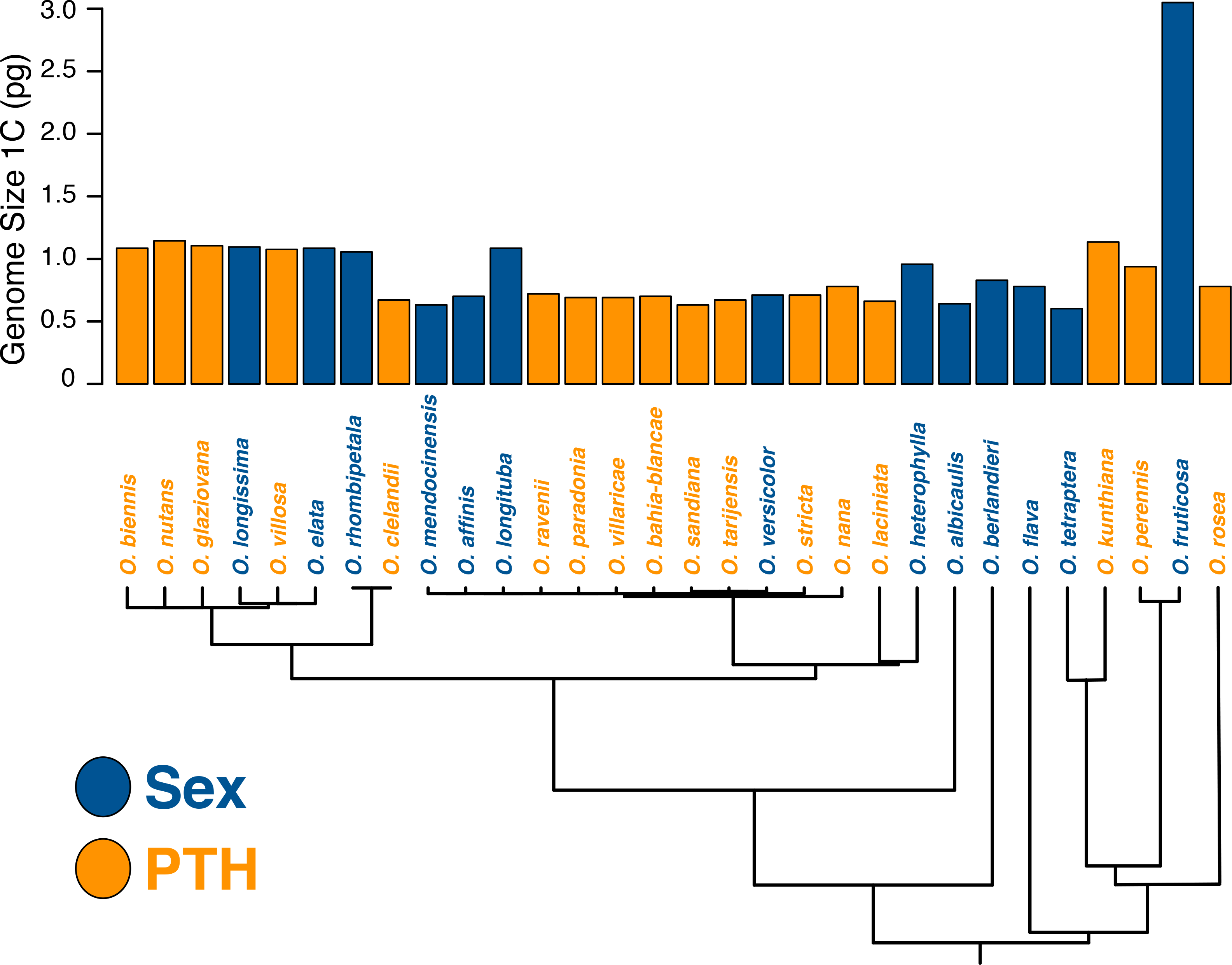
Molecular phylogeny of *Oenothera* used in the PGLS analysis. Haploid genome size estimates (pg; 1 pg = 978 Mb) are plotted on top. Note, *O. fruticosa* is a polyploid and not included in the statistical analyses.

### Phylogenetic analysis

To control for statistical non-independence due to shared evolutionary history (Felsenstein 1985), we accounted for phylogeny in our statistical analysis. We inferred the phylogeny of the 30 species using the previously generated phylogeny of *Oenothera* in Johnson et al. (2009). Briefly, we sequenced two plastid (*trnL-trnF* and *rps16*) and three nuclear gene regions (*PgiC*, *ITS*, and *ETS*) from 121 species and created a maximum likelihood phylogeny using RAxML 7.0.4 (Stamatakis 2006). The tree was made ultrametric using non-parametric rate smoothing in TreeEdit (http://tree.bio.ed.ac.uk/software/treeedit) and then pruned to include only the 30 species studied here. For further details of the phylogeny see Johnson et al. (2009).

To start, we tested whether the data exhibited significant phylogenetic signal using Pagel’s λ (Pagel 1999) implemented in the phylosig function in the *phytools* package (Revell 2012) of R version 3.0.2 (R Development Core Team 2013). This package assesses the significance of phylogenetic signal by performing a likelihood ratio test against the null hypothesis that λ = 0. Next, we performed phylogenetic generalized least squares (PGLS; Butler and King 2004) regression between reproductive system (sex was coded as 0 and PTH as 1) and genome size using the *ape* (Paradis et al. 2004) and *geiger* (Harmon et al. 2008) packages in R. We performed the PGLS tests under both neutral (Brownian motion) and stabilizing selection (Ornstein-Uhlenbeck) models of trait evolution. Akaike’s Information Criterion (AIC) was used to determine which model best described the data.

### DNA isolation and Illumina sequencing

Seeds of *O. elata* (sex) and *O. biennis* (PTH), two species with relatively large genome sizes, and of *O. villaricae* (PTH), one of relatively small genome size (Supplementary Table 1), were imbibed at 4°C overnight in the dark with 3% Plant Preservative Mixture (Plant Cell Technology Store, Washington, DC, USA). Seeds were then sown on the substrate surface and grown until the early rosette stage (for details see Greiner and Köhl, 2014). For DNA isolation, 50 mg of leaf material was frozen in liquid N_2_ and ground using a mixer mill. After adding 775 µl of IGEPAL-buffer [1.9% IGEPAL, 1.9% CTAB, 130 mM Tris/HCl (pH 8.0), 130 mM EDTA (pH 8.0), 1.3% PVP-40, 1.8 M NaCl, 130 mM B(OH)_3_], 125 µl 1-thioglycerol, and 100 µl RNase A (50 mg/ml), samples were incubated at 60°C for 30 minutes under medium shaking. Cell debris was removed by centrifugation. The supernatant was treated twice with chloroform/isoamylalcohol (24:1) and subsequently with phenol/chloroform/isoamylalcohol (25:24:1, pH 7.5). The phenol/chloroform/isoamylalcohol treatment was repeated until the aqueous phase was no longer “reddish”. DNA was precipitated with 1/10 volume of 5 M NH_4_-acetate and 1 volume of isopropanol (−20°C), incubated at −20°C for 15 min, collected by centrifugation, and washed with 70% and/100% EtOH (−20° C). Resolved DNA was further purified using the Genomic DNA Clean & ConcentratorTM kit (Zymo Research Cooperation, Irvine, CA, USA). DNA was sequenced at the Max Planck-Genome-Centre Cologne (Germany) on an Illumina HiSeq2500 platform, 100 bp paired-end, utilizing a TruSeq DNA library (375 bp insert size). All sequences will be deposited to the Sequence Read Archive (http://www.ncbi.nlm.nih.gov/sra).

### Repetitive content analysis

To assess whether variation in genome size could be attributed to differential accumulation of repetitive elements such as TEs, we determined the repetitive content in three species of varying genome size and reproductive system. To characterize repeats we ran RepeatExplorer (Novak et al. 2013), a *de novo* graph-clustering pipeline for repeat characterization (Novak et al. 2010), which is implemented in the Galaxy platform (http://galaxyproject.org). We filtered reads for quality, keeping only reads with a Phred quality score of at least 20 over 90% of their length. RepeatExplorer joins reads together in clusters based on sequence similarity and then matches these clusters against RepBase (Jurka et al. 2005) to identify repeats. We ran the pipeline on one sample per species under default settings (Table 1).

**Table 1.** Clustering statistics for Repeat Explorer

|  | *O. villaricae* | *O. elata* | *O. biennis* |
| --- | --- | --- | --- |
| Total number of reads used in analysis | 1001571 | 1012981 | 729978 |
| Number of reads in clusters | 813712 | 857862 | 639924 |
| Number of clusters | 24496 | 19474 | 12197 |

## Results

### Variation in genome size

We detected almost two-fold variation in genome size among the diploid species surveyed (Figure 1; Supplementary Table 2). Estimated genome size ranged from 0.64 pg in *O. mendocinensis* (sex) and *O. sandiana* (PTH), to 1.16 pg in *O. nutans* (PTH). The South American clade (Subsection *Munzia*: *O. mendocinensis*, *O. nana*, etc.) consistently had the smallest genomes, whereas the recently radiated North American *O. biennis* clade (Subsection *Oenothera*) consistently had the largest genome sizes. The so-called “B clade” (sensu Wagner et al. 2007; e.g. *O. flava*, *O. tetraptera*, *O. perennis*, *O. fruticosa*, *O. rosea*, *O. kunthiana*) varied the most in genome size, reflecting polyploidy (exact ploidy level unknown) in one species (*O. fruticosa*) and deep divergence among multiple lineages. Excluding the polyploid *O. fruticosa* (sex), the mean genome size was 0.85 +/−0.036 s.e. pg. C-value estimates will be submitted to Kew Garden’s Plant DNA C-Value Database (http://data.kew.org/cvalues/)

### Phylogenetic signal

We detected significant phylogenetic signal in genome size across the species examined. Pagel’s λ was 0.74 (*P* = 0.0141 that λ >0) suggesting that phylogenetic non-independence should be taken into account in statistical analyses.

### Phylogenetic generalized least squares analysis

We found no significant relationship between sexual reproduction and genome size. The lack of an effect of sex on genome size holds regardless of whether we assume that genome size evolves under a neutral Brownian motion model (df = 29*, P* = 0.828) or moving towards a selective optimum in an Ornstein-Uhlenbeck’s model of stabilizing selection (df = 29, *P* = 0.8162). Comparing the AIC scores suggests that the Brownian motion model (AIC = −103.4203) better describes the data than the Ornstein-Uhlenbeck model (AIC = −42.8108).

### Repetitive content

Repetitive elements were abundant in all three species examined. After filtering for quality, about 80% of all reads formed clusters (Table 1). In all three species TEs made up most of the repetitive content, with the dominant TEs being long terminal repeat *gypsy* and *copia* elements (Figure 2). In all three species, we estimated that TEs make up ∼ 35-40% of the genome.

**FIGURE 2.**
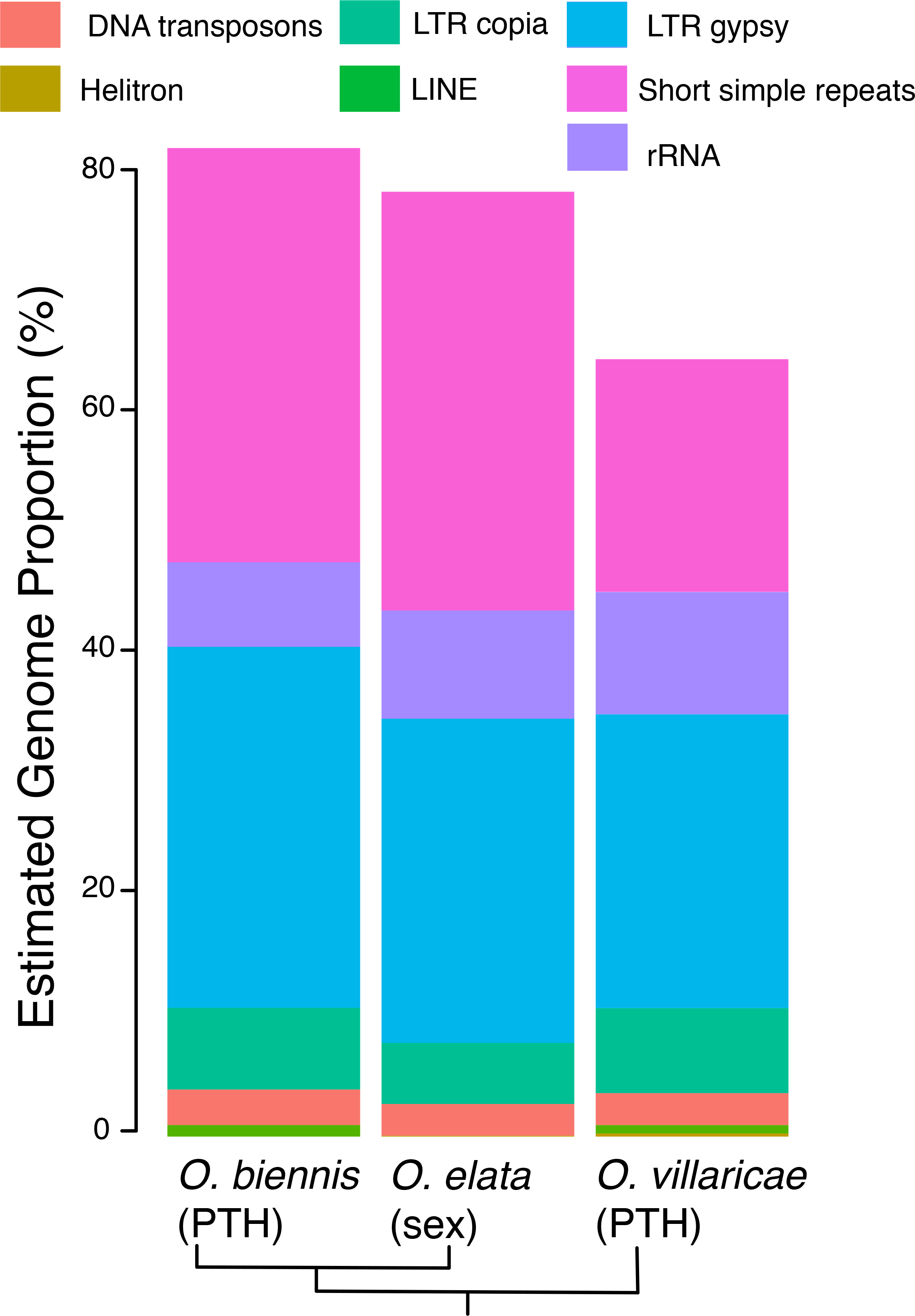
Repetitive content of three *Oenothera* species as estimated by RepeatExplorer (Novak et al. 2013), including two species of relatively large genome size (*O. elata* and *O. biennis*) and one of relatively small genome size (*O. villaricae*).

To investigate whether the genomes differed in other kinds of repeats, we combined the sequences annotated as “simple repeat”, “satellite”, and “low complexity” under the label “short simple repeats”. These repeats made up a larger proportion in the relatively large genomes *O. elata* and *O. biennis* (∼ 35%), than in the smaller *O. villaricae* (∼ 20%; Figure 2) and this difference was statistically significant (Pearson’s chi-square test of independence, χ^2^ = 59360.79, df = 2, *P* = 2.2 × 10^−16^).

## Discussion

Studies of genome size variation have a long history (Mirsky and Ris 1951). References to this variation have featured heavily in the arguments about the role of non-selective processes in the evolution of genome complexity (Lynch and Conery 2003; Lynch 2007; Whitney and Garland 2010; Whitney et al. 2010; Lynch 2011; Whitney et al. 2011) and more recently in the debate associated with the ENCODE Project Consortium’s claim that 80% of the human genome can be assigned a biochemical function (ENCODE Project Consortium 2012; Eddy 2012; Graur et al. 2013; Doolittle 2013; Kellis et al. 2014; Palazzo and Gregory 2014). With estimates from some 7,000 species and a 2,400-fold variation in genome size (Leitch and Leitch 2013), studies of plant genomes have much to contribute to these and other debates.

Here, we presented genome size estimates in thirty species in the evening primrose genus *Oenothera* and found no evidence that sex explains the almost two-fold variation in genome size. Instead, evolution of genome size was fairly conserved within *Oenothera* and best explained by neutral genetic drift, as opposed to a model of stabilizing selection towards an optimum, or a model that ignores evolutionary history. Moreover, contrary to the reasoning outlined in the Introduction, we found no evidence that genome size variation in *Oenothera* can be attributed to transposable element abundance. Instead the observable difference in genome size appeared to be due to accumulation of short simple repeats.

Central to the hypothesis that sexual species should have larger genomes than asexual relatives is that sexual reproduction should facilitate the spread of transposable elements (Hickey 1982; Charlesworth and Langley 1986; Morgan 2001; Dolgin and Charlesworth 2006). Our results are in line with two recent studies highlighting that differences in TE abundance are not the only reason why lineages may differ in genome size. For example, in *Eucalyptus*, a member of the same order as *Oenothera* (Myrtales), a difference in abundance of tandem repeats rather than TEs is responsible for the 110 Mb difference in genome size between the closely related *E. grandis* and *E. globulus* (Myburg et al. 2014). In *Arabidopsis*, the highly selfing *A. thaliana* has fewer TEs and smaller genome than its outcrossing relative *A. lyrata* (Hu et al. 2011), which is likely due to accumulation in the outcrosser rather than a loss of TEs in the selfer (Slotte et al. 2013), consistent with TE transmission advantage in an outcrossing lineage. However, genome size variation within *A. thaliana* does not seem be due to TEs. In particular, Long et al. (2013) detected a 10% variation in genome size among 180 Swedish *Arabidopsis thaliana* lines, which was due to differential accumulation of 45s rDNA rather than TEs. What determines whether genome size difference will be due to TEs or simple repeats remains unclear.

Although there is evidence from multiple systems that sex may promote the spread of TEs (Zeyl et al. 1996; Arkhipova and Meselson 2000; Schaack et al. 2010), there is also abundant evidence that asexuality is associated with a reduction in the efficacy of selection (reviewed in for example Glemin and Galtier 2012). Consistent with this, a recent study of 13 sexual and 16 PTH *Oenothera* species found evidence of a reduction in the efficacy of selection in PTH species (Hollister et al. submitted). If the magnitude of this reduction in the efficacy of selection is large enough, it may dilute the transmission advantage associated with sex. This kind of dilution effect could be magnified if the sexual species vary in selfing rate. However, variation in selection efficacy due to variation in selfing rate is unlikely to explain our results because all sexual species examined, except *O. versicolor* and *O. mendocinensis*, are either self-incompatible or highly herkogamous, i.e. having spatially separated anthers and stigma.

The results from our within-genus comparison corroborate those of the multi-family study of the role of outcrossing rate in genome size evolution by Whitney et al. (2010). In their paper, they find no effect of outrcossing rate on genome size and suggest that an effect of mating system in their analysis could have been obscured by rapid mating system shifts. In our analysis, the short time scale due the recent divergence between species may mean that any effect of reproductive system have yet to materialize. Using computer simulations to examine changes in TE copy number following a shift to asexual reproduction, Docking et al. (2006) found that it took around 50,000 generations to reach a new equilibrium. During the first 30,000 generations following the shift to asexuality, copy numbers often increased dramatically, but even with low levels of excisions all elements were lost. At any given point in time, however, variation among lineages was very high. Thus although quantitative predictions from computer simulations will be sensitive to specific parameter values assumed, they do highlight that sampling a limited number of lineages could fail to detect a correlation between sex and TE copy number, especially if divergence times are small such is the case in *Oenothera*. If the rapid shifts of mating and reproductive systems mean that the effect on TE and genome size evolution on both short (within genus) or long (multi-familiy) time scales is difficult to capture, a possible middle road could be to examine within family variation. The plant family Brassicaceae, home of *Arabidopsis*, is a good candidate for such a family. Within the Brassicaceae, genome size varies 16-fold across the almost two hundred species (Lysak et al. 2009; Johnston et al. 2005), species vary extensively in mating system (Vekemans et al. 2014), and the large number of sequenced reference genomes (Haudry et al. 2013) make it suitable for comparative genomic analysis.

Here, we have performed the first comprehensive study of the role of sex in genome size evolution in plants. Whereas it has long been clear that sexual reproduction affects various aspects of genome evolution, including in the evening primroses (Ellstrand and Levin 1980; Hersch-Green et al. 2012; Hollister et al. submitted), it appears to have played no role in genome size evolution in *Oenothera*.

## Acknowledgements

We thank Niroshini Epitawalage and Victor Mollov for assistance in the glasshouse, Brian Husband and Paul Kron for discussion and advice on flow cytometry, and Elena Ulbricht-Jones and Liliya Yaneva-Roder for providing the DNA samples for Illumina sequencing. Liliya Yaneva-Roder is further acknowledged for excellent technical assistance during the establishment of the phenol-based DNA isolation protocol for *Oenothera*. Seeds were generously donated by USDA GRIN, Algonquin Park Visitor Centre staff, S.C.H. Barrett, Ornamental Plant Germplasm Centre (OPGC), J. Meurer, W. Wagner, and R. Raguso. J.A.Å is supported by a Junior Fellowship from Massey College, S.G. by the Max Planck Society and the Deutsche Forschungsgemeinschaft (DFG), M.T.J.J. and S.I.W. by the Natural Sciences and Engineering Research Council (NSERC). We declare no conflict of interest.

